# Gene expression models based on a reference laboratory strain are bad predictors of *Mycobacterium tuberculosis* complex transcriptional diversity

**DOI:** 10.1101/091082

**Authors:** Alvaro Chiner-Oms, Fernando González-Candelas, Iñaki Comas

## Abstract

Species of the *Mycobacterium tuberculosis* complex (MTBC) kill more people every year than any other infectious disease. As a consequence of its global distribution and parallel evolution with the human host the bacteria is not genetically homogeneous. The observed genetic heterogeneity has relevance at different phenotypic levels, from gene expression to epidemiological dynamics. However current systems biology datasets have focused in the laboratory reference strain H37Rv. By using large expression datasets testing the role of almost two hundred transcription factors, we have constructed computational models to grab the expression dynamics of *Mycobacterium tuberculosis* H37Rv genes. However, we have found that many of those transcription factors are deleted or likely dysfunctional across strains of the MTBC. In accordance, we failed to predict expression changes in strains with a different genetic background when compared with experimental data. The results highlight the importance of designing systems biology approaches that take into account the tubercle bacilli, or any other pathogen, genetic diversity if we want to identify universal targets for vaccines, diagnostics and treatments.

## INTRODUCTION

Tuberculosis (TB) has killed more persons in the last two hundred years than any other infectious disease. Today, TB is still the main infectious disease agent in the world, accounting for about 1,4 million deaths plus 390,000 associated to HIV co-infection (WHO | Global tuberculosis report 2016, 2016). In humans, the disease is caused by *Mycobacterium tuberculosis* and *Mycobacterium africanum*, which belong to the *Mycobacterium tuberculosis* complex (MTBC) along with other species that cause the disease in animals. The bacterium infects the host through the respiratory track. Once in the lungs, it is phagocyted by macrophages which typically are encapsulated in a granuloma (Orme & Basaraba, 2014). The bacteria can be dormant and survive inside the granuloma during months, years or even decades, in an asymptomatic disease state called latency (Getahun *et al*, 2015). The transition from latency to an active disease state depends on biological features of the bacteria, the host, environmental factors and the interactions among all of them (Comas & Gagneux, 2009). These interactions are not completely understood yet (Kondratieva *et al*, 2014). Moreover, animal models are widely used but they do not reproduce perfectly the human-pathogen interaction (Vilaplana & Cardona, 2014; Orme & Basaraba, 2014).

One way to approach the complexity of the host-pathogen-environment triangle is through systems biology. In the case of TB, systems biology approaches have produced encouraging results in the identification of persistence genes (Dutta *et al*, 2014), the pharmacokinetics and pharmacodynamics of TB drugs inside the granuloma (Pienaar *et al*, 2015; Lalande *et al*, 2016) and the identification of drug resistance mechanisms (Peterson *et al*, 2016). Overexpression experiments as well as chromatin-immune-precipitation sequencing (ChIP-Seq) data has been used to produce a detailed map of the interactions and regulatory logics of more than 200 transcription fators (TFs) in H37Rv, the laboratory reference strain (Rustad *et al*, 2014; Minch *et al*, 2015). The enormous quantity of data generated is publicly available and can be used to study the regulatory interactions of the bacteria in several ways (Turkarslan *et al*, 2015).

However, little attention has been paid to the fact that H37Rv is a clinical strain used in laboratories for decades and that for many aspects does not represent the whole species. Therefore, it is expected that natural perturbations in the inferred H37Rv biological networks introduced by naturally occurring mutations in clinical strains will change the gene model architecture and predictions and the underlying regulatory network. In fact, along their evolutionary story *M. tuberculosis* complex strains found in humans have diverged in 7 different lineages (Comas *et al*, 2013). The aforementioned H37Rv strain belongs to lineage 4. Most of the genetic variation among lineages in *M. tuberculosis* results from wide genomic deletions and point mutations (Gagneux *et al*, 2006). It is also known that the maximum genetic distance between strains of different lineages is 2,188 SNPs (Coscolla & Gagneux, 2014). The phenotypic role of mutations defining lineages has been extensively studied and some of them are clearly linked to transcriptional differences between the MTBC lineages (Rose *et al*, 2013; Dinan *et al*, 2014; Homolka *et al*, 2010). It is also clear that one single mutation affecting regulatory processes can impact dramatically on the virulence of the pathogen (Gonzalo-Asensio *et al*, 2014; Pérez *et al*, 2001). In fact, a novel live vaccine attenuated carrying a deletion in the key regulator PhoP is currently on phase 1B of clinical trials (Spertini *et al*, 2015). In addition, several studies have shown that bacterial genetic diversity has epidemiological implications and genetic differences among lineages lead to differences in the immune response and disease progression in the host (Portevin *et al*, 2011; de Jong *et al*, 2008; Coscolla & Gagneux, 2014; Gagneux *et al*, 2006; Reiling *et al*, 2013). As a result, novel diagnostics, vaccines and treatments maybe compromised by failing to account for the circulating diversity as has recently been described for several diagnostics tests based on the detection of the protein of *mpt64* (Ofori-Anyinam *et al*, 2016).

Thus, we are completely blind as of whether the topology of the regulatory networks available and the gene expression mathematical models derived from H37Rv can be extrapolated or not to other strains of the MTBC and how the regulatory modulations are affected by the existing bacterial diversity. In this work we derive new gene expression models by pooling existing H37Rv data and explore their predictive power on genome wide expression patterns when natural variations (mutations) seen in clinical strains are introduced. We show how different experimental set-ups can affect the inferred models of gene expression and regulatory influence and how far we are from predicting only from transcriptomic data the impact of genetic polymorphisms at a genome-wide expression level.

## RESULTS

### Building and validation of gene expression models based on lineage 4 H37Rv strain

Taking advantage of recently published experimental datasets testing the regulatory influences of known TFs, we defined gene expression models for the laboratory reference strain H37Rv. The datasets included transcription factors overexpression experiments (TFOE) for ~200 TFs (~700 microarray experimental tests) (Rustad *et al*, 2014). In addition, for each TF we have also used information about the binding sites of transcription factors along the *M. tuberculosis* H37Rv genome (Minch *et al*, 2015). Firstly, we used the data to model the expression behavior of each gene in H37Rv by determining the number of TFs (regressors) influencing the expression and secondly, we established the dynamics of this impact by assigning a coefficient that modulates gene expression changes. Finally, we test whether the observed influence is exerted directly or not by overlapping transcription and transcription factors binding sites information.

Gene expression models were built using a backward step-wise algorithm (Figure 1, see Material and Methods for details). The approach generated 3,960 putative gene expression models. When, in addition, we required evidence of physical interaction the number of initial putative models was reduced to 755. Therefore, our putative gene models accounted for 98.3% of the coding capacity of the genome when physical interaction was not required, and 19.24% of the coding capacity of the H37Rv genome when we used the ChIP-Seq data. Secondly, we cross-validated all models in the two datasets and then compared them with random models to discard spurious results. Following this approach, we discarded 2,744 models for the TFOE and retained 1,216 (30.8%). For the case of ChIP-Seq data, only 29 models were retained (3.74% of the initial models) (Figure 2A). The models derived from TFOEs alone included a larger number of TFs per model, as expected due to the larger number of regulatory events incorporated. On the contrary, the models derived from the combination of TFOEs and ChIP-Seq data had fewer TFs influencing the expression as they only include those TF physically bound to the gene (Figure 2B). In summary, our approach shows the relevance of performing serial statistical validations of expression models derived from experimental data. The structure of the final models is included in the expanded view (File EV1).

**Figure 1.**
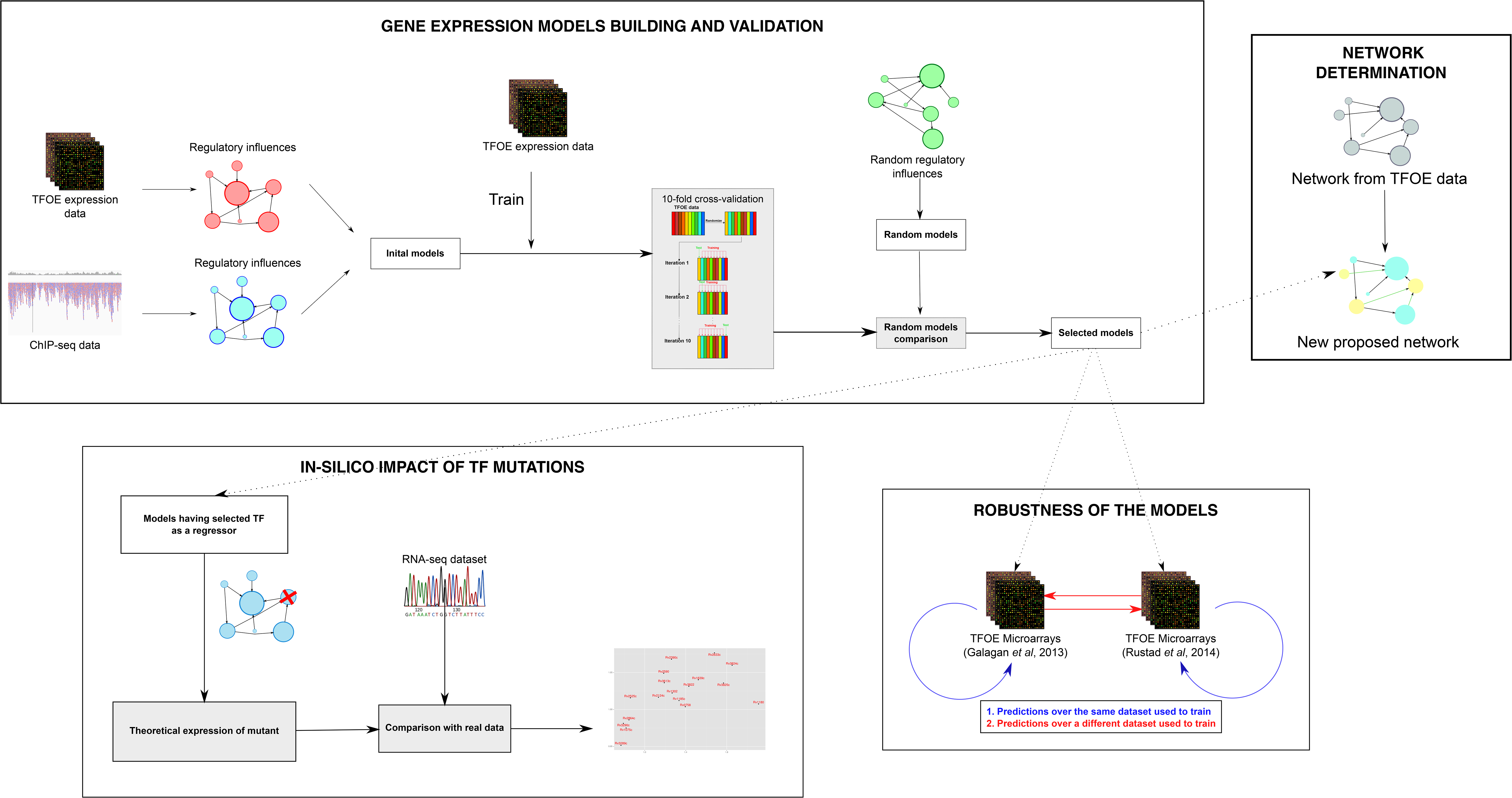
Workflow to build and validate gene expression computational models. Initial models were derived from the regulatory relationships derived from the TFOE and ChIP-Seq data. They were trained with dataset from Rudstadt *et al*. 2014 (for more details about the different datasets used, check Materials and Methods). These initial models went through a 10-fold cross-validation process, to discard low accuracy models. The remaining ones were compared with random models. Those showing better performance than the random models were selected as the final models. These selected models were used to (i) check the robustness of the models (ii) derive a new regulatory network (iii) *in-silico* predict the impact of a TF deletion.

**Figure 2.**
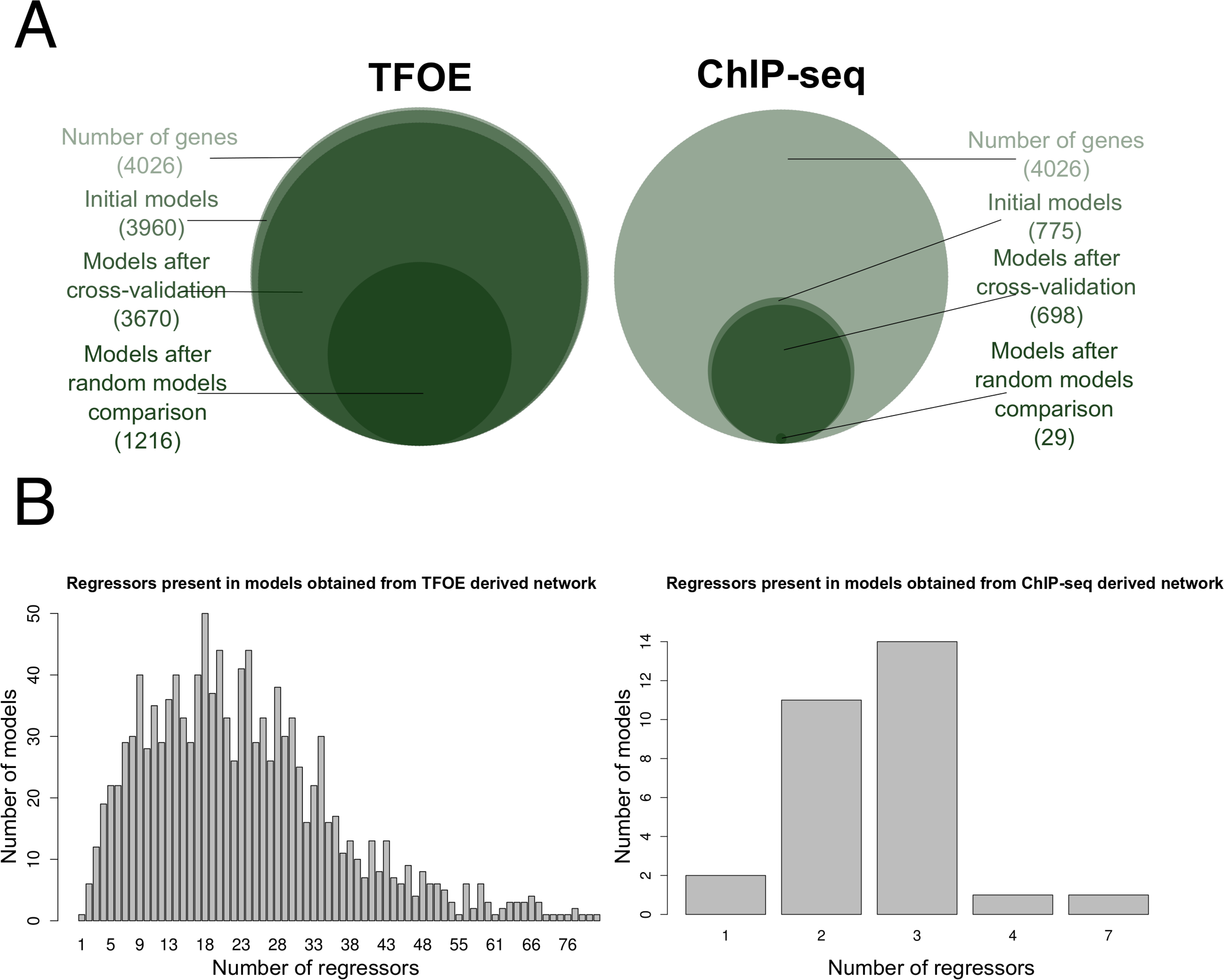
Gene expression computational models. A Overview of the results obtained during the building process and refining steps of the computational models derived from the TFOE data (left) and the ChIP-Seq data (right). B Distribution of the number of TFs affecting the target gene on each network derived model.

During the building of predictive models, we assigned coefficients to each regressor (TF) impacting gene expression by using TFOE data as a training set (Rustad *et al*, 2014). To evaluate how robust the prediction was to noise, we compared the predictions with the expression values obtained in a previous, analogous TFOE experiment (Galagan *et al*, 2013). In the case of the TFOE-derived models, we were able to predict the expression values of the corresponding gene in Galagan *et al*. (2013) only for 128 genes (10.52% of gene models, pFDR ≤ 0.01). In the models incorporating physical interactions, only 10 genes (34.48% of gene models) were observed with no significant differences between measured and predicted expression values, with a pFDR ≤ 0.01. In fact, a mere comparison of average expression values for each gene across the two datasets (Galagan *et al*. vs Rustad *et al*.) already shows that experimental noise has a great influence. Mean expression values for the same gene were not statistically significant in only 18 cases (pFDR < 0.01, Figure EV1). This result already suggests the presence of substantial experimental noise in otherwise analogous experiments.

### Regulatory network based on statistically validated interactions

The 1,216 expression models obtained from the TFOE dataset include 11,253 regulatory relationships. These relationships are the ones selected after applying the backward step-wise method in the models building process (see Material and Methods for details). Although all of them lead to a lower Bayesian Information Criterion in their respective models, most of these relationships are based on a weak regulatory signal. To select for the strongest links between TFs and gene regulation influence we kept those leading to significant gene expression (two-fold change) according to the TFOE data (Rustad *et al*, 2014). With these subsets of regulatory relationships, we built a new regulatory network. The new network (Figure 3A) comprises 3,396 regulatory events across 1,102 genes (37.15% of the events and 38.76% of the genes from the network proposed by Rustad *et al*.). The main centrality statistics comparing the initial and the inferred network are shown in Table EV1. The distribution of the in-degree parameter of the network (Figure 3B) reveals that most genes are regulated by an intermediate number of factors whereas a minority is regulated by a large or small number of them. On the other hand, the distribution of the out-degree parameter follows the expected power-law distribution (Junker & Schreiber, 2008) with most TFs regulating a small amount of genes and a few genes affecting the regulation of many.

**Figure 3.**
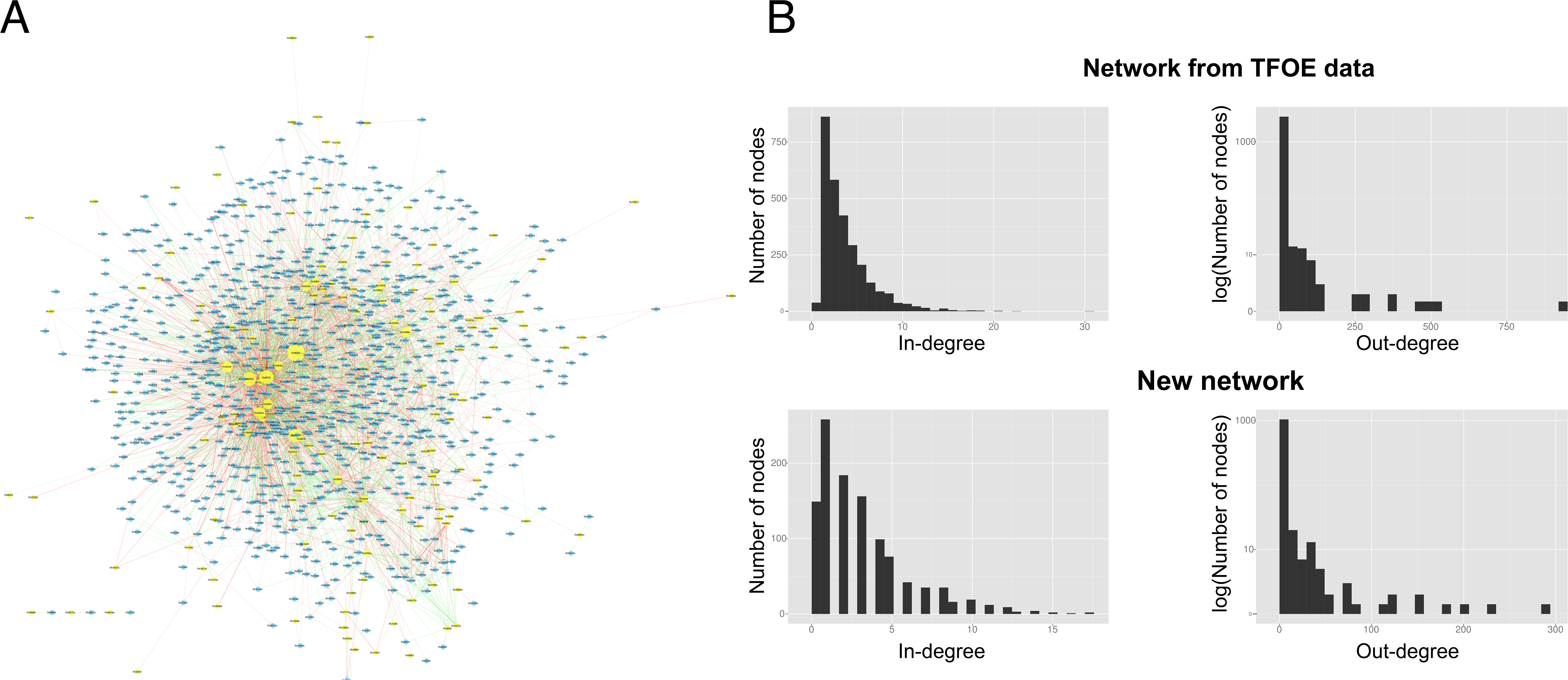
Validated gene expression network. A New regulatory network proposed. Yellow nodes represent genes that code for transcription factors while blue nodes represent target genes for the corresponding TF. Sizes of the nodes are proportional to the number of genes regulated by them. Green edges indicate positive regulations while red edges indicate negative regulation. The transparency of the edges is related with the TF → gen influence (the darker the edge, the higher the regulatory effect). B Comparison of the out- and in-degree distributions between the network derived from TFOE data and the new proposed network.

In agreement with the original networks of Rustad *et al*. and Galagan *et al*., in this new regulatory network Rv0023 and Rv0081 are the TFs that regulate the largest number of genes (672 and 627, respectively). Thus, they are regulatory hubs of *M. tuberculosis*. On the other hand, the gene Rv3202c is the one with the largest number of TFs influencing its expression, as it is indirectly regulated by 26 TFs. This gene has ATPase and helicase activities (Lew *et al*, 2011). The regulatory subnetwork of Rv3202c is related to regulatory DNA and RNA processes, as well as response to external stimuli, transport and secretion. The network containing the regulatory links statistically validated in this analysis can be found in File EV2 and can be downloaded from https://tbgenomics.wordpress.com/resources/.

### Transcription factors are not universally conserved in the MTBC

Once we had available gene expression models and a regulatory network for H37Rv we tried to predict the phenotypic effect of natural genetic variation observed in circulating clinical strains. To do so, we first examined the degree of conservation of the studied TFs across the *Mycobacterium tuberculosis* complex. Previous studies have identified mutations in the genes that code for the PhoPR system in MTBC strains that had important effects on pathogen’s virulence (Gonzalo-Asensio *et al*, 2014), so that not only SNPs in the regulatory regions of the TF but also those located in the coding region could lead to differences in the TF activity. Thus, we focused our analyses on mutations falling in regulatory regions but also on those coding mutations that might impair the normal function of the TF.

Using a collection of genome-wide SNPs and indels from a large number of clinical strains (Comas *et al*, 2013), we have identified a total of 28 transcription factors (TFs), among those present in the TFOE data (Rustad *et al*, 2014), that are missing or likely dysfunctional (as defined in Material and Methods) in one or more clinical strains including four affecting complete lineages of the *M. tuberculosis* complex (Table EV2 and Table EV3) (Figure 4). Some of these transcription factors are deleted in complete lineages and sublineages as they are in known regions of difference (RD) used as phylogenetic markers (Gagneux *et al*, 2006). For example, Rv1994c and Rv2478c are in RD743 and RD715 and they affect the entire lineage 5 (Mostowy *et al*, 2004). Those lineages represent up to 50% of tuberculosis cases in West Africa (Gehre *et al*, 2016). We have also identified single point mutations disrupting the normal functioning of some TFs. This is the case for *sirR* (Rv2788). An early stop codon mutation was found in all the strains of lineage 1. In the proposed regulatory network, Rv2788 regulates 22 genes (Figure 5B), in accordance 16 of those genes are differentially expressed in lineage 1 strains with respect H37Rv using available RNAseq data (Rose *et al*, 2013). In our estimates (Figure 5A), lineage 1 accounts for roughly 18% of the strains causing active tuberculosis cases each year (almost 1.9 million cases/year), hence the relevance of taking into account the circulating genetic diversity when building comprehensive regulatory networks.

**Figure 4.**
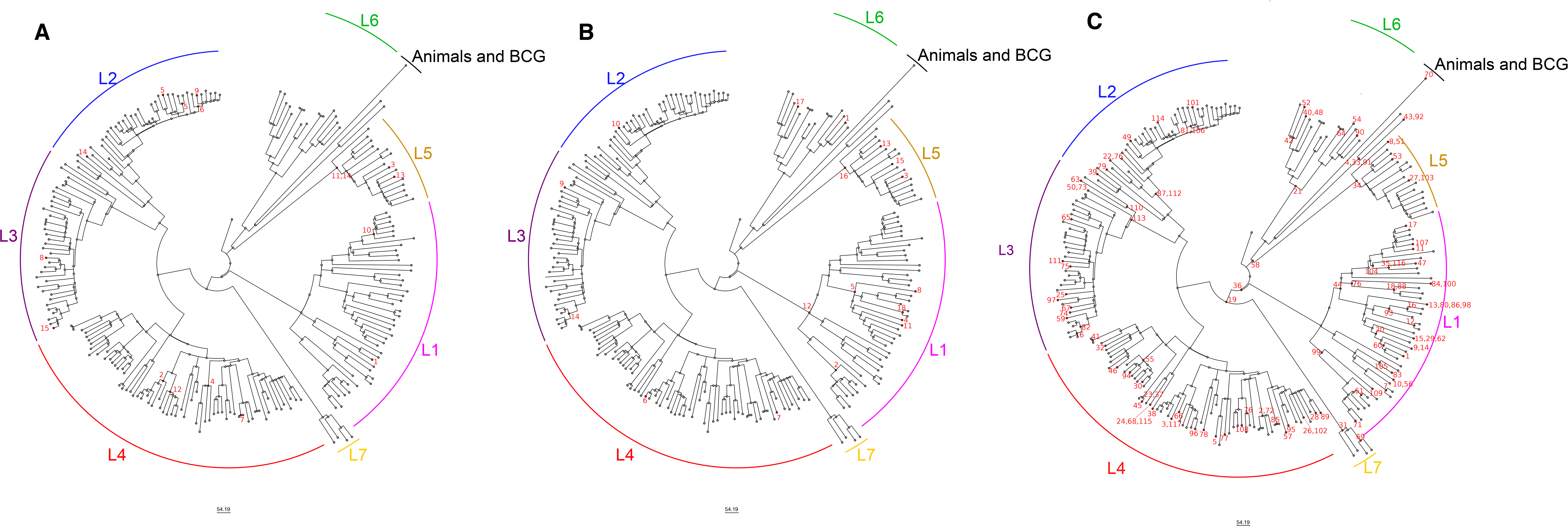
MTBC phylogeny comprising the seven major lineages. The figure represents the number of TFs missing in one or more clinical strains of the MTBC from Comas et al. (2013). Each panel shows the same phylogeny and the mutations affecting a TF are mapped to the corresponding branch in the tree and highlighted in red. Label numbers correspond to entries in Table EV2, Table EV3 and Table EV4. The mutations considered are either partial or complete deletions of the TF (A), single point mutations leading to gain or loss of stop codons (B) and single point mutations affecting the regulatory region of a TF (C).

**Figure 5.**
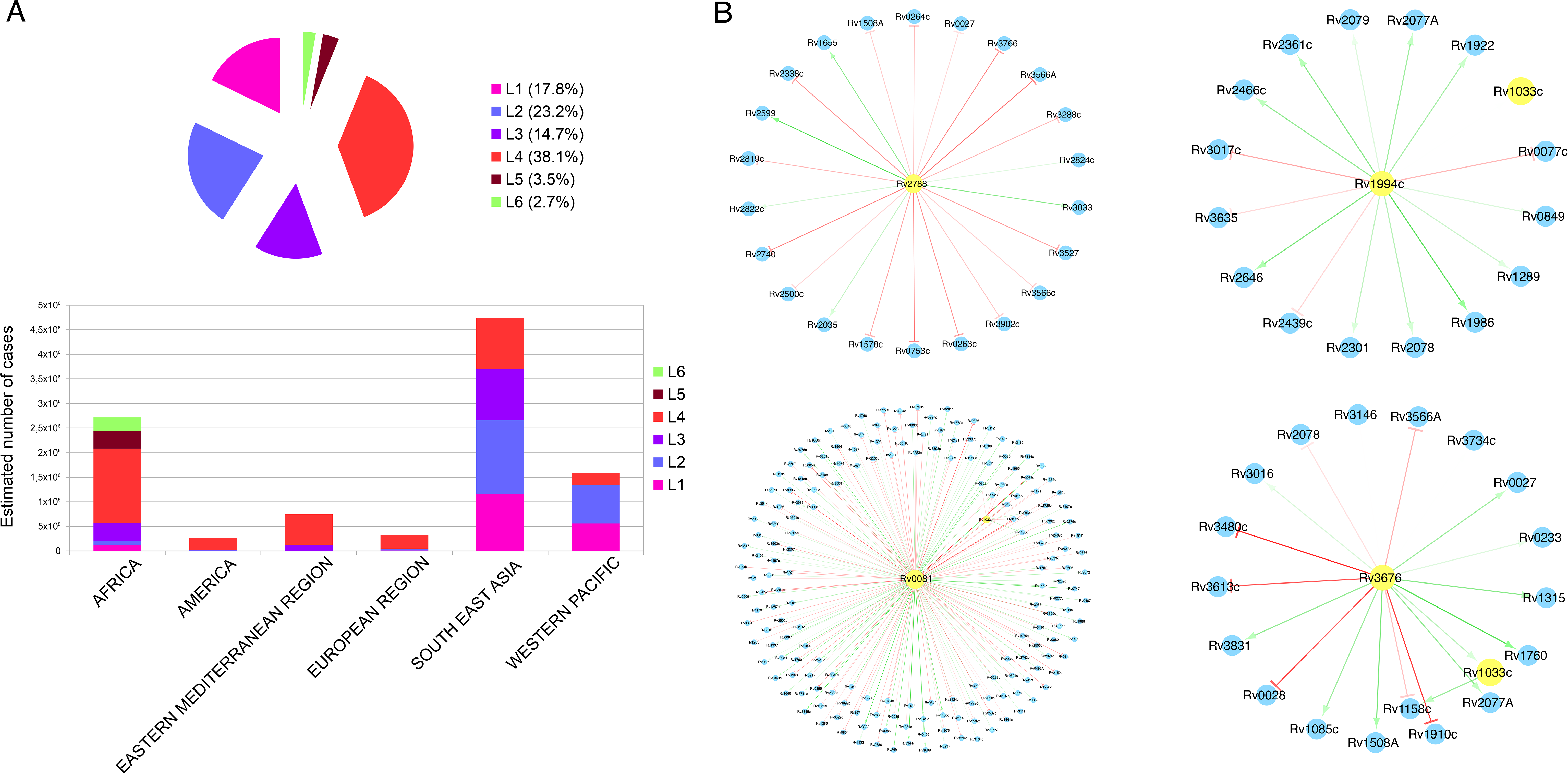
Global incidence of the different lineages and representative examples of mutations affecting complete lineages. A Pie chart showing the estimated number of annual tuberculosis cases attributed to each lineage and barplot showing the incidence of the different lineages by region. Lineage 7 is not shown due to its low incidence in global terms (Comas *et al*, 2015). The data related to the disease incidence by region come from the WHO (WHO | Global tuberculosis report 2016, 2016) and the lineage abundance for each region from a previous work (Gagneux *et al*, 2006). B Examples of regulatory subnetworks of transcription factors affected by mutations in one or more lineages. From upper-left to lower-right: regulatory subnetwork of Rv2788 (early stop-codon in all lineage 1 strains); regulatory subnetwork of Rv1994c (deleted in all lineage 5 strains); regulatory sub-network of Rv0081 (SNP in regulatory region found in all the strains screened from lineage 2,3,4 and 7) and regulatory sub-network of Rv3676 (SNP in regulatory region in all the strains from lineage 3).

Next, in order to check the main biological processes involved we analyzed the relative abundance of Gene Ontology (GO) terms in the regulatory subnetworks for each affected TF (Table EV2 and Table EV3, expanded view File EV3). Most of the TFs identified as missing in clinical strains have an important role, with a direct or indirect regulatory influence in up to 210 genes. The GO analysis showed that the affected TFs are involved in a wide range of processes, related with metabolism, regulation, pathogenicity and response to external stimuli. Some deletions affecting TFs appear in single strains, such as one affecting Rv1994c in a strain of lineage 2 or Rv1776c in a strain of lineage 3. A deletion of gene Rv1985c, a known antigen, was also found in a group of strains belonging to lineage 1. It is also remarkable that a stop-codon gain mutation was found in Rv0465c (also known as *ramB*) in one strain of lineage 4. RamB is related to the glyoxilate cycle in the pathogen and it has been proposed to play an important role in the adaptive response of the bacteria to different host environments during infection (Micklinghoff *et al*, 2009). Moreover, the regulatory subnetwork of *ramB* is involved in several processes such as regulation of RNA biosynthesis, response to hypoxia or interaction with the host.

We have also identified 117 SNPs located in the regulatory regions of 44 TFs (Table EV4, Figure 4). Most of these SNPs affect primary or alternative transcription start sites (TSS) as defined in a previously published work (Cortes *et al*, 2013); two of them correspond to antisense TSS and two more were internal TSS. Seventy-four of these SNPs affect one single strain, with the remaining 43 affecting more than one. Interestingly, only a few of them impact complete lineages, such as T89200G, which impacts the master regulator Rv0081 in modern lineages 2, 3, 4 and 7 (76% of the circulating strains), or C422745T, which impacts Rv0353 in all lineages except 5 and 6. Rv0081 regulates 188 genes (including *tcrR*, which also regulates 26 genes) (Figure 5B) so a SNP potentially affecting its regulation could have an important effect on the regulatory network of the bacteria (Gonzalo-Asensio *et al*, 2014). Besides, we have found one homoplasic SNP (C2965900T, which affects Rv2642) that have emerged independently in strains of 3 different lineages. It has been shown previously that some of the SNPs screened are already known as affecting the expression of their corresponding TF (Table EV4). For example, SNP G3500149A has been reported to be involved in the regulation of TF Rv3133c in Beijing strains (lineage 2), as it creates a -10-box leading to the overexpression of the DosR regulon (Rose *et al*, 2013; Domenech *et al*, 2016).

### *In-silico* expression prediction of genetic backgrounds observed in a clinical and in a vaccine strain

To explore how well the H37Rv-based expression models and the validated network predict the impact of the genetic background in the transcriptional landscape of the bacteria we have selected a lineage 1 strain (T83) from the comparative genomics analysis. For T83 there is publicly available expression data set (Rose *et al*, 2013) and we have identified a deletion in TF Rv1985c (Table EV2) and an early stop-codon in Rv2788 (Table EV3). By reducing the expression of Rv1985c and Rv2788 to its minimum level, we created gene models mimicking the T83 genetic background. With these modifications, we were able to identify significant differences in the predicted expression values in 148 of the 169 genes that have Rv1985c or Rv2788 as regressors (pFDR < 0.05). We compared the predictions with available experimental data from a RNA-seq dataset. To explore how well models perform qualitatively we tested if we could predict the direction of the gene expression change (induction or repression). The predictions agree with the experimental data for 71 genes but failed for 77. A Cohen’s kappa test was performed and no agreement was found between the predicted and the real values (kappa = -0.06) (Table EV5). We performed also a quantitative test so we compared the expression values of the 148 predictive models showing significant differential expression with those genes in which we observed an adj-pvalue < 0.05 in the differential expression analysis between T83 and H37Rv. Sixty-four coincidences were found. There was no correlation between the predicted vs the real fold-change in the expression of the 64 genes identified (Pearson correlation coefficient = 0.08, p-value = 0.48). Although conclusions from a single strain are necessarily provisional, it is also true that the mutations in T83 are present in several strains of lineage 1 (Table EV2 and Table EV3, Figure 4). Thus, from the limited data available we speculate that gene expression models based on H37Rv and derived from TFOE are not likely to predict accurately enough the transcriptional landscape of *M. tuberculosis* complex lineages.

Once was not possible to predict expression changes in strains other than H37Rv, we tested the impact of knocking out a single TF in the H37Rv genetic background. As an example, we selected PhoP for several reasons: (i) it is one of the main regulators in MTBC (Pérez *et al*, 2001; Galagan *et al*, 2013); (ii) it is the main gene deleted in a vaccine candidate that is already in clinical trial phase 1B (Spertini *et al*, 2015); (iii) there are large datasets published of the expression changes in knockout strains using two different approaches, microarray (Gonzalo-Asensio *et al*, 2008) and RNAseq (Solans *et al*, 2014); and (iv) there is strong evidence that mutations in the PhoPR regulatory regions impact fitness of clinical strains on the human host (Gonzalo-Asensio *et al*, 2014).

From the TFOE, we identified 218 models in which *phoP* (Rv0757) is present as a regressor. When the expression value of *phoP* was lowered to the minimum level allowed by the model, thus simulating that the gene has been knocked-out, there were 188 genes in which statistical significant differences between mutant and normal expression values were observed (pFDR < 0.05). To contrast these predictions with experimental data we used the expression differences between isogenic clinical strains with or without the *phoP* knock-out mutation. Among the 188 gene models influenced by PhoP, 10 of them are in a list of 78 genes influenced by PhoP according to (Gonzalo-Asensio *et al*, 2008). The regulatory influence in these 10 cases, derived from predictions and compared with the experimental values of Gonzalo-Asensio *et al*. is shown in Table EV6.

We also contrasted our predictions with an RNA-seq dataset of a *phoP* knockout H37Rv strain (Solans *et al*, 2014). Again, we first compared if the predicted expression follows the same direction as the ones from the RNA-seq dataset. In 96 cases the predictions agree with the experimental values but in 92 cases the predictions failed (Table EV6). The Cohen’s kappa test shows a slight agreement between the real and the predicted values (kappa = 0.05). After that, we compared the 188 predictive models showing differential expression with the genes showing differential expression in the dataset (adjusted p-value < 0.05) and we found only 9 coincidences. For these 9 genes in common, we obtained almost no correlation (Pearson correlation coefficient = 0.59, p-value = 0.09) between the predicted and measured gene expression fold-change in the mutant (Figure 6). To test whether the regulatory influence is exerted through direct interaction or indirect influence, we used the ChIP-Seq data from the physical binding network. Only one model included *phoP* as a regressor, Rv2590. This result suggests that PhoP acts indirectly over many genes, as shown previously (Solans *et al*, 2014; Galagan *et al*, 2013).

**Figure 6.**
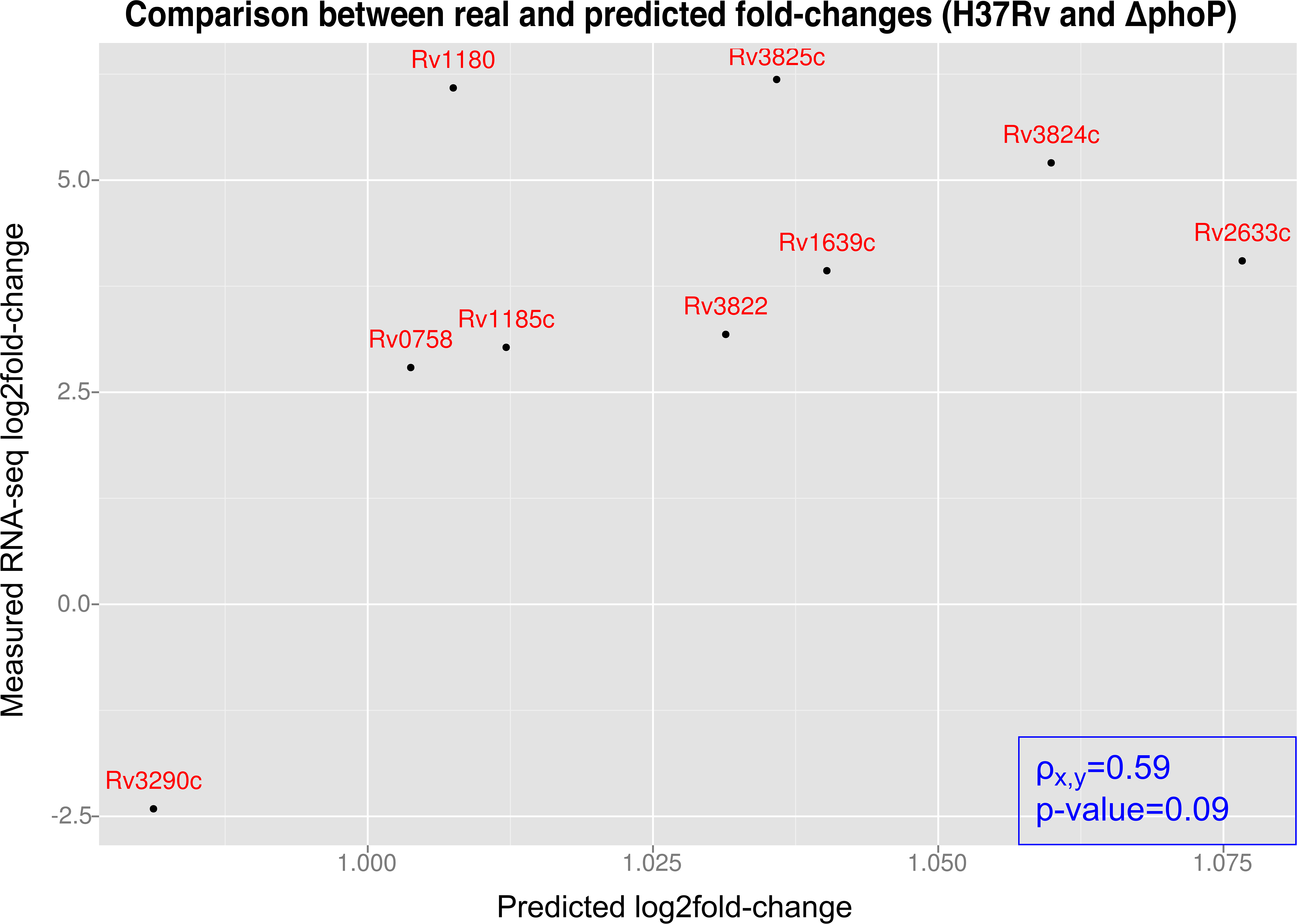
Comparison between experimental and predicted fold-changes. The y-axis corresponds to the measured log2 fold-change in gene expression between the WT strain and the *Δphop* strain in Solans *et al*. (2014) The x-axis corresponds to the predicted fold-changes calculated with the predictive models developed in this work.

Next, we tested whether the lack of correspondence between our predictions and the experimental data might be due to the former being obtained from TFOE whereas the later were defined after analyzing a knock-out strain. Figure 7A shows a graphical comparison between the ChIP-Seq coverage of the over-expressed, the knock-out mutant, and the wild-type strains. Using the wildtype coverage vs the *phoP* mutant coverage as a negative control, we are able to infer the binding sites of PhoP in the H37Rv strain (Solans *et al*, 2014). By comparing these results with the binding sites inferred from the TFOE strains, we observed differences in several genes (Figure 7B). For example, in the Rv1101 gene there is no evidence of PhoP regulation when the mutant and the wildtype are compared. However, a peak appears in the overexpressed strain and strong regulatory evidence has been reported (Rustad *et al*, 2014). In total, from 139 genes predicted to be regulated by PhoP from the TFOE data and 51 from the mutant data, only 16 genes overlap. Thus, different methodologies to test the function of a gene (overexpression versus deletion) partially account for the limited predictive power of the H37Rv gene expression models even when the mutant is derived from a H37Rv background.

**Figure 7.**
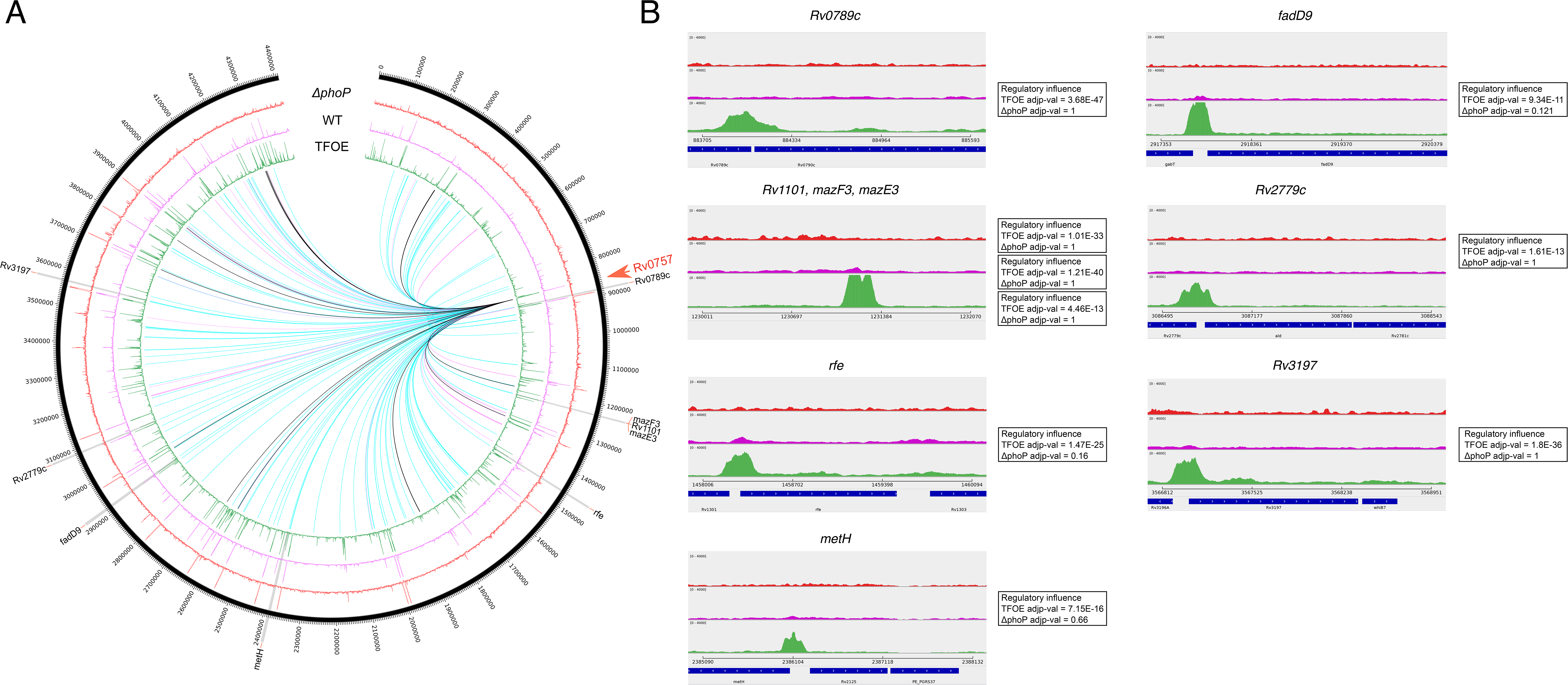
ChIP-Seq coverage comparison and regulatory influences between the *phoP* knockout, wild-type and *phoP* overexpressed strains. A The outer circle represents genomic positions. From outside to inside: ChIP-Seq coverage of *phoP* mutant, wild-type and TFOE. The inner links represent the regulatory influence of *phoP* derived from TFOE (blue), mutant strain (purple) and the overlap between them (black). B Detail of genes with regulatory influences derived from TFOE that do not match with the evidence from WT and the mutant strain.

## DISCUSSION

Systems biology approaches rarely accommodate information about natural polymorphisms in the systems studied. However, these polymorphisms can contribute to new genetic backgrounds with important consequences at the phenotypic level. Evolutionary systems biology has developed as a discipline that aims at generating quantitative insight on complex systems to model their evolution. One of its main focus is to predict the effect of mutations on phenotypes. However, this kind of approach has been rarely applied to non-model organisms and even within model systems rarely to more than one genetic lineage (Gasch *et al*, 2016). Thus, the actual power of systems biology models in predicting the effect of new mutations on non-model system still has to be evaluated (Tagu *et al*, 2014). This is the case for tuberculosis, in which both the pathogen and the host are not genetically homogeneous and genetic variation in any of them may impact disease progression and outcome (Comas & Gagneux, 2011). Therefore, systems biology-derived models must accommodate the potential impact of host and bacterial genetic heterogeneity in order to make universal predictions in a pathogen of global distribution.

We have tested the predictive power of state-of-the-art *M. tuberculosis* regulatory networks and expression models when the system is disturbed by (i) several mutations associated to a clinical strain with a genetic background different to the training dataset, and (ii) a knock-out mutation in the key regulator PhoP in the reference strain used for the training dataset. Both for the genetic background and single mutations predictions our results show very little overlap between the genes predicted to be significantly impacted and those experimentally determined. Overall, these results suggest that our predictive models only grab a minor part of the true phenotypic variation.

### Experimental noise impacts gene expression models prediction

One striking result is that our gene expression models in most cases are not statistically different than randomly generated models and that they do not correctly predict the expression level even in analogous TFOEs experiments (Galagan *et al*, 2013). Several biological factors might explain the discrepancies between predicted and observed expression values. First of all, our results show that the statistical validation of gene expression models is essential to tease out methodology-dependent effects that may or may not correspond to actual biological differences and contrasted biological effects. Only 1,216 gene models derived from the TFOE dataset and 29 from the ChIP-Seq were significantly different from randomly-generated gene expression models. It is well known that in some cases noise is introduced by subtle expression differences between cells and can have a biological role (Macneil & Walhout, 2011; Chalancon *et al*, 2012). In fact, a recent study with *Bacillus subtillis* has demonstrated that gene expression noise contributes to phenotypic heterogeneity, providing some advantage in certain environments (Mugler *et al*, 2016). However, in most cases it just represents perturbations introduced by differences in the experimental setting (Parekh *et al*, 2016; Grün *et al*, 2014). This suggests that in the future when trying to understand the impact of different perturbations in the system, such as genetic mutations, the noise introduced by the experimental setting must be taken into account, especially in genes with a low expression level (Malone & Oliver, 2011).

Secondly, the regulatory network analyzed is highly dependent on the experimental methodology used. Overexpression of transcription factors is a common, widely used technique to identify regulatory influences but it can fail to make accurate predictions when an increase in gene expression has no physiological effect or, on the contrary, overestimate the regulatory effect due to a loss of specificity (Blais & Dynlacht, 2005). A recent work with *Mycoplasma pneumoniae* demonstrated that the overexpression of regulatory molecules (asRNAs in this case) leads to an overestimation of the regulatory effect of these molecules (Lloréns-Rico *et al*, 2016). Also, the ChIP-Seq technique could introduce false positives when overexpressing the TF (Park *et al*, 2013).

Furthermore, knock-out mutant approaches are universally used to understand the role of a gene but the comparison with overexpression approaches may not be accurate. There are several examples in our data in which genes whose expression is clearly impacted in *phoP* knock-out experiments show no impact in TFs overexpression datasets. On the other hand, these regulatory networks take into count only the relationships inferred from transcription factors. It is known that regulatory mechanisms of *M. tuberculosis* are more complex. In some cases, increasing the expression of a TF does not lead to an increase of its activity, as some TFs need to be phosphorilated or undergo conformational changes in different conditions to be activated. Increasing the amount of the TF but not changing the conditions could make its activity undetectable. Besides, other studies (Arnvig *et al*, 2011) have shown that there is a large number of sRNAs in bacteria that regulate genic expression. Their effect is not reflected in this type of networks and analyses, as we only look for the amount of mRNA produced by a gene. Finally, the amount of mRNA does not always agree with the amount of translated protein (Maier *et al*, 2009; Vogel & Marcotte, 2012). However, new experimental techniques such as CRISPRi are being tested in *M. tuberculosis* and several other organisms to characterize and modify gene expression (Singh *et al*, 2016). The introduction of CRISPR-based techniques in this research field could allow overcoming some of the disadvantages of current techniques and to define regulatory influences more accurately.

### Current models are bad predictors of MTBC transcriptional diversity

Given the low concordance between predictions and expression datasets generated with the same methodology, we also expected low predictability when we tried to mimic the impact of a different genetic background. Given the well-known impact of mutations on transcriptional activities, we first carried out a comparative genomics analysis of the conservation of known TFs in the MTBC. Our analyses show how the different TFs tested in H37Rv are not universally conserved. Some of those mutations (either deletions or single point mutations) impact complete lineages and up to 76% of the circulating strains.

In our case we have predicted the transcriptional landscape of a lineage 1 strain. We have found 64 coincidences between the genes predicted to be impacted and those found in an RNA-seq experiments (Rose *et al*, 2013). Strain T83 belongs to lineage 1 and its genetic distance to H37Rv, the strain used to build the models, is more than 1,800 SNPs (Comas *et al*, 2013). Thus, other genetic differences besides those found in TFs between this strain and the one used to infer the regulatory influences will certainly impact the genome-wide transcriptional landscape of T83. For example, we have mapped 2 SNPs in the regulatory region of TF Rv0353 (Table EV4) in T83. The potential influence of these SNPs is not covered by the current models as well as those of other regulatory layers that possibly differ between lineages. In addition, we have shown before that specific SNPs of lineage 1 alter the expression levels of sense and antisense transcripts by means of the appearance of new TSSs (Rose *et al*, 2013). Accordingly, we found a better agreement between predicted and observed expression changes when we introduced the effect of deleting a single TF (PhoP) in a H37Rv background.

In summary, current models of regulatory networks cannot be used to accurately predict the impact of genetic variation on expression levels of the regulated genes. Noise as well as missing regulatory layers also represent a major obstacle for modelling expression changes. This might be the main reason explaining why, of the initially calculated models, only 30.8% from the TFOE derived models and 3.74% in the ChIP-Seq models showed a better performance than random generated models. To overcome these limitations, systems biology approaches including protein-protein interaction networks, *in-silico* modeling, metabolic flux analyses, or transposon-based functional characterizations are being carried out (Peterson *et al*, 2014; Ma *et al*, 2015; Garay *et al*, 2015; Crouser, 2016; Padiadpu *et al*, 2016).

From this work it is also clear that there is a need for experimental data from representative samples comprising the natural diversity of the *Mycobacterium tuberculosis* complex. This is a major effort if we take into account that the data used for this work and coming from H37Rv alone was derived from almost 200 TFs genetic constructs and 700 microarray experiments. Generating comprehensive models for all major human and animal lineages of the *M. tuberculosis* complex will represent a challenge in the years to come. In addition, multiomics data is also desirable in order to capture the major regulatory layers (Monk *et al*, 2016). The challenge will be to integrate all of them in a manner that can inform each other (Ma *et al*, 2015) and to accommodate and predict the role of existing human and bacterial genetic diversity (Comas & Gagneux, 2011). This a major obstacle for using systems biology approaches in *M. tuberculosis* complex to prioritize targets for biomedical research.

## MATERIALS AND METHODS

### Datasets and techniques used

The main microarray expression datasets were obtained from Rustad *et al*. (2014) (available at GEO with accession numbers GSE59086). Also, the ChIP-Seq data was obtained from Minch *et al*. (2015). The TFOE derived network used to compare with the TFOE network generated in this work was obtained from Rustad *et al*. (2014) (available at http://networks.systemsbiology.net/mtb/). The *phoP* mutant data was obtained from Solans *et al*. (2014) (available at GEO under accession number GSE54241). The RNA-seq data from lineage 1 (including the T83 strain) was obtained from Rose *et al*. (2013) (available at EBI ENA under accession number ERP002122). The H37Rv RNA-seq data was obtained from Arnvig *et al*. (2011).

The R statistical language was used to perform all the analyses (Team, 2015), mainly the Bioconductor set of packages (Huber *et al*, 2015). All the methodology is summarized in Figure 1.

### Model construction

The regulatory relations used for model construction were obtained from the Rustad *et al*. (2014) dataset downloaded from the MTB network portal (http://networks.systemsbiology.net/mtb/). We selected all the regulatory interactions with adjusted p-value <= 0.01 regardless the fold-change in the expression values. In consequence, all the statistically significant regulatory influences (even the weak ones) were taken into account. From Minsch *et al*. (2015) we selected the physical bindings which demonstrated regulatory effect over the gene expression of the target.

We used a backward step-wise methodology for constructing the computational models. The following process was performed for each target gene in the TFOE and the ChIP-Seq derived models.

1. All the TFs affecting the gene were selected as regressors for the model. Besides, the RNA polymerase alpha chain gene, *rpoA* (Rv3457c), and the sigma factor gene, *sigA* (Rv2703), were included as normalization factors. Also, the interactions between TFs were taken into account. The model structure (Galagan *et al*, 2013) was:

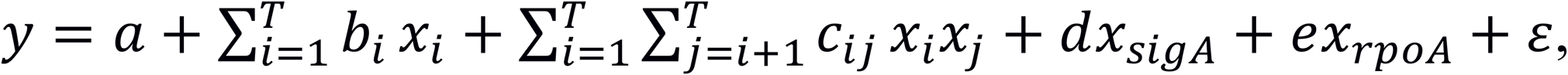

where *y* is the target gene expression, *x*_*i*_ the expression values of the selected TFs (from *i*=1 to *T*), *a, b,d* and *e* the linear coefficients in the regression model, *c* are the interaction coefficients, and *ɛ* is the error term.

2. A linear regression model with all the TFs selected as regressors and based on the previous structure was constructed. Later, the model was parameterized using microarray data from Rustad *et al*. (2014).

3. The Bayesian Information Criterion (BIC) and the Akaike Information Criterion (AIC) associated to the model were calculated. To limit the overfitting error we used the BIC in the TFOE derived models because it penalizes models with a large number of regressors (Faraway, 2009). In turn, the AIC was used when calculating models from the ChIP-Seq dataset given the low number of regressors involved.

4. We sequentially eliminated the regressor whose removal from the model led to the highest decrease of the BIC/AIC. The remaining TFs were selected and we returned to step 2. In case we did not observe a decrease in the BIC/AIC after the removal of any regressor, we considered that model as optimal for the corresponding gene.

5. A Fisher’s F-test was performed to check the null hypothesis that the retained regressors do not have predictive power (Faraway, 2009). P-values were adjusted to multiple testing by Benjamini and Hochberg false discovery rate (FDR)(Benjamini & Hochberg, 1995) and all models with adjusted p-value >= 0.05 were rejected.

### Cross-validation of models

We checked the initial models obtained above in a 10-fold cross-validation.

For each gene:

1.- The optimal model selected was parameterized using a random subset of the 90% TFOE dataset as a training-set.

2.- After that, the remaining 10% of the dataset was used as a test-set to make predictions. A Fisher F-test was performed to check differences between residuals of the training-set and the test-set. Also, the Root Mean Squared Error (RMSE) (Chai & Draxler, 2014) when predicting over the testset was obtained.

3.- Steps 1 and 2 were repeated 10 times (10-fold cross-validation)

In this case, we retained the models that showed no differences between predictions over the training-set and the test-set, by comparing the average adjusted p-value of the F-test over the 10 iterations (α >= 0.05). In some cases, we could not find differences between residuals but the squared error was high. In consequence, we also rejected the models with RMSE > Q3 + 1.5 * IQR, as they will be considered as outliers of the RMSE distribution (Tukey, 1977).

### Comparisons to random models

We considered each TF of the datasets as a potential regulator. For each gene, we listed all the TFs that do not have a regulatory influence over it. From this list we created 100 random subsets of TFs. The number of elements in each subset was equal to the number of real factors with regulatory influence over the corresponding gene. With this random subset of TFs, we followed the steps described above to create the random models. Also the 10-fold cross-validation was performed for each random model.

For each model, a Welch’s t-test to compare the distribution of p-values from the 10-fold cross-validation of the real model versus the random ones was performed. P-values from Welch tests were adjusted by Storey’s method (Storey, 2003). Tests showing a pFDR <= 0.01 were accepted as having a better fit than random models. Also, the RMSE distributions of random models versus the real ones were tested by means of a Welch’s t-test, correcting the p-values with Storey’s method. Tests showing a pFDR <= 0.05 were accepted.

### Evolutionary conservation of TFs within the MTBC

To determine whether TFs have been conserved along the evolutionary history of the MTBC we analyzed an available dataset of natural polymorphisms in 219 representative strains of the complex, encompassing all known lineages and geographic distribution of the species (Comas *et al*, 2013). A custom script was used to search for TFs that have at least 25% of their gene length deleted. A manual inspection was performed over every TF deletion found to filter false positives and mapping errors. Also, single nucleotide polymorphisms (SNPs) leading to stop-codon gains or losses, and point mutations affecting any TF regulatory regions (Cortes *et al*, 2013) were extracted from this dataset. The terminology used to classify the TSSs follows Cortes *et al*. (2013).

We defined the regulatory sub-network associated to each TF as that defined by the one-step distance nodes to the TFOE network. To understand the potential role in major physiological functions, for each sub-network we studied the enrichment in certain functional categories as defined by the GO classification (The Gene Ontology Consortium, 2014). The Gene Set Enrichment (GSE) analysis was carried out using a combination of the BINGO tool (Maere *et al*, 2005) and the Cytoscape software for visualization (Shannon *et al*, 2003). The enrichment test identifies the most abundant GO terms in the sub-network compared to all the possible terms present in the complete annotation, using a hypergeometric test (sampling without replacement).

### RNA-seq analysis

Expression data from H37Rv and lineage 1 strains was obtained from Rose *et al*. (2013). The differential expression analysis was performed using DESeq2 package (Love *et al*, 2014).

Differentially expressed genes were those with an adjusted p-value of 0.05. Differential expression analysis was performed over H37Rv versus all lineage 1 strains to check the differences in those genes regulated by Rv2788 (*sirR*). In addition, we specifically analyzed differentially expressed genes between between T83 (lineage 1) and H37Rv (lineage 4).

### Predicting the impact of genetic polymorphisms

To predict the impact of the genetic background on the transcriptional landscape of the *M. tuberculosis* complex we chose a lineage 1 strain, namely T83. As a result of the analyses of conservation degree of the TFs described above, we have identified two genetic mutations in lineage 1 which likely have a major impact on the functionality of TFs. One of the mutations corresponds to a deletion affecting TF Rv1985c while the other is in an early stop codon in TF Rv2788 (Table EV2 and Table EV3). To simulate a transcriptional landscape for lineage 1, the expression values of both TFs were set to the minimum value found in the training dataset as standard regression models approaches requires to make predictions in the same data range to that used to parameterize the model (Faraway, 2009). The T83 gene expression predicted values were compared to those obtained from H37Rv expression models. To evaluate the differences between predicted and real values we performed a Welch’s t-test over the expression values. P-values were adjusted by Storey’s method. A pFDR < 0.05 was considered for accepting the difference between models as significant. For the genes that showed differential expression, we calculated the log2 fold-change between H37Rv and T83. We compared these values with those obtained from the RNA-seq analysis shown above. For a qualitative approach, we checked if the changes in the expression values follow the same direction (positive for induction and negative for repression). We constructed a 2x2 matrix with the predicted effect vs the measured effect and a Cohen’s kappa test was performed over this matrix to check the agreement between prediction and real data. After that, we wanted to check the quantitative accuracy of the models so we selected the genes that showed differential expression in both, the RNA-seq dataset (adjusted p-value < 0.05) and in the predicted expression to calculate the Pearson correlation.

Similarly, to make predictions on a *phoP* mutant in a H37Rv background we set its expression value to the minimum value found in the training dataset for this TF. The analysis was performed following the steps described above. To analyze how accurate, the models reflect fold-changes in experimental data we used an RNA-seq data from a *phoP* mutant and H37Rv (Solans et al, 2014) as explained in the previous section. We compared the log2-based fold-changes between the predictive models and experimental data comparisons.

To compare the ChIP-Seq coverages in the different cases, we obtained raw data from the wild-type strain and the *phoP* mutant from Solans *et al*. (2014). We also downloaded ChIP-Seq data from the overexpression experiment of *phoP* (Rv0757_B167) from the MTB network portal (http://networks.systemsbiology.net/mtb). The circular diagram was constructed with the Circos tool (Krzywinski *et al*, 2009) and the values from the regulatory influence were extracted from the TFOE dataset.

## ACKNOWLEDGEMENTS

This work was funded by projects BFU2014-58656-R from Ministerio de Economía y Competitividad (Spanish Government) and PROMETEO/2016/122 from Generalitat Valenciana (to FGC), Ministerio de Economía y Competitividad (Spanish Government) research grant SAF2013-43521-R, and the European Research Council (ERC) (638553-TB-ACCELERATE) (to IC). ACO is recipient of a FPU fellowship from Ministerio de Educación y Ciencia FPU13/00913 (Spanish Government)

## AUTHORS’ CONTRIBUTIONS

IC. Conceived the work. IC, FGC, ACO designed the experiments. ACO carried out the experiments. IC, FGC, ACO analyzed the data and wrote the manuscript.

## FIGURE LEGENDS

**Figure EV1. Evaluation of accuracy and comparison of model’s behavior between different datasets.**

To evaluate the robustness of the complete models to changing experimental conditions we compared the models built with the expression values observed in an analogous experiment (Galagan *et al*, 2013). The main goal of the models is to make predictions under different conditions and with several data sources. Therefore, apart from training the models with the same dataset used to calculate the models and the regulatory networks (Rustad *et al*, 2014) we trained them with the other analogous dataset. In this figure, dataset A is the one obtained from Rustad *et al*, (2014) and dataset B from Galagan *et al*, (2013).

(A) Root mean squared error comparison (RMSE) for the models obtained from TFOE data. Values when training and testing with dataset B (green), training and testing with dataset A (blue), training with dataset A and testing with dataset B (red) and training with dataset B and testing with dataset A (yellow).

(B) Plot showing RMSE values for TFOE derived models. Index refers to the list of models sorted by RMSE. The green arrows mark those models having no differences between predicted and measured mean expression. The left plot shows the case of training with dataset A and testing with dataset B while the right plot shows the reverse case. In the left plot, 128 genes show no differences between real and predicted values in terms of equaliy of means while in the right plot 33 genes show no statistical differences (pFDR ≤ 0.01).

(C) RMSE comparison for the models obtained from ChIP-Seq data. Values when training and testing with dataset B (green), training and testing with dataset A (blue), training with dataset A and testing with dataset B (red) and training with dataset B and testing with dataset A (yellow).

(D) Plot showing RMSE values for ChIP-Seq derived models. Index refers to the list of models sorted by RMSE. The green arrows mark those models having no differences between predicted and measured mean expression. The left plot shows the case of training with dataset B and testing with dataset A while the right plot shows the reverse case. In the left plot, 10 genes show no differences between real and predicted values in terms of equality of means while in the right plot only 3 genes show no statistical differences (pFDR ≤ 0.01).

